# Stimulation of otolith irregular fibers produces a rostro-caudal gradient in activity in the vestibular nuclear complex (VNC), but not the vestibulocerebellum (VeCb)

**DOI:** 10.1101/2024.09.20.614117

**Authors:** Avril Genene Holt, Shana King, Danial Naqvi, Rod D. Braun, David S. Bauer, Marie Anderson, William Michael King

## Abstract

The vestibular system is important for posture, balance, motor control, and spatial orientation. Each of the vestibular end organs have specialized neuroepithelia with both regular and irregular afferents. In otolith organs, the utricle and saccule, afferents most responsive to linear jerk (jerk - derivative of acceleration) are located in the striola and project centrally to the vestibular nuclear complex (VNC) as well as the uvula and nodulus of the vestibulocerebellum (VeCb). The pattern of central neuronal activation attributed to otolith irregular afferents is relatively unknown. To address this gap, c-Fos was used as a marker of neuronal activity to map the distribution of active neurons throughout the rostro-caudal extent of the VNC and VeCb. Immunohistochemistry for c-Fos was performed to assess activation of VNC and VeCb neurons in response to a linear jerk stimulus delivered in the naso-occipital plane. Activated neurons were distributed throughout the VNC, including the lateral vestibular nucleus (LVe), magnocellular medial vestibular nucleus (MVeMC), parvocellular medial vestibular nucleus (MVePC), spinal vestibular nucleus (SpVe), and superior vestibular nucleus (SuVe). Notably, after stimulation, the MVePC exhibited the greatest number of c-Fos labeled nuclei. Significant increases in c-Fos labeling were found in mid-rostrocaudal and caudal regions of the VNC in the LVe, MVe, and SpVe. Additionally, c-Fos labeling was observed across all regions of the VeCb after jerk stimulation. Significant increases in the number of labeled nuclei were found throughout the rostro-caudal extent of the nodulus and uvula. However, jerk stimulated increases in activity for the paraflocculus were restricted to the caudal VeCb. The distribution of neuronal activity suggests that regions receiving the greatest direct otolith input exhibit the most substantial changes in response to otolith derived, irregular fiber stimulation.

**Highlights:**

- Nuclei with descending projections (LVe, MVePC, and SpVe) demonstrated the greatest change in activity after naso-occipital jerk stimulation.
- Naso-occipital jerk stimulation preferentially activates caudal VNC neurons
- Naso-occipital jerk stimulation activates neurons throughout the VeCb
- Jerk stimulation in the naso-occipital plane has the greatest effects on activity in VNC and VeCb regions with the greatest inputs from afferents originating in gravity receptors

## Introduction

The vestibular system is essential for posture, coordinated movement, and spatial orientation. The vestibular system works with the visual system to maintain balance by stabilizing gaze during head and body movement. In mammals, including rats, head movement is detected in the periphery by two otolith organs and three semicircular canals on either side of the head. These organs transduce linear and angular acceleration in response to changes in head position relative to gravity or head rotations in space respectively. These signals are sent to the central nervous system to allow quick adjustments of postural musculature to maintain an upright position or stabilize the body axes during locomotion.

The otolith end organs, the utricle and saccule, contain specialized regions called maculae, within which lie the sensory epithelia. These sensory regions contain hair cells that synapse with afferents of the vestibular nerve (cranial nerve VIII). The afferent inputs are arranged such that afferents responsive to high linear acceleration (calyx only afferents) are located in a specialized region called the striola. Within the central nervous system, calyx-only afferents do not appear to be confined to one region, but project to all four subdivisions of the vestibular nuclear complex (VNC) as well as the uvula and nodulus of the vestibulocerebellum (VeCb) [1–3]. Since calyx only fibers receiving signals emanating from otoliths are particularly responsive to abrupt linear acceleration, rapid head jerks have been used as robust stimuli to produce synchronous discharges in utricular and saccular irregular fibers in rodents. These synchronous discharges are detected as far field potentials called vestibular short latency evoked potentials (VsEPs) and provide a quantitative estimate of peripheral and central synchronous irregular afferent activity [4–6]. Of the 6-8 peaks, analysis of VsEPs has primarily focused on the earliest peaks, P1, P2, and 3, which are believed to represent the synchronous discharge of irregular afferents [4, 7–9]. More recently, VsEP waves P1 and P2 have been used to assess the effects of high intensity noise stimulation (e.g. noise) on the periphery [10–13]. While additional experiments still need to be performed, current studies suggest that P2 and P3 have central components [7].

There are few studies that address activation of the vestibular periphery in response to jerk stimulation of otolith organs [14] and even fewer studies examining the effects of jerk on central responses [15]. The projections of afferents from the vestibular periphery to the VNC and VeCb are known to be broadly distributed across subdivisions [16–19], but central patterns of activity resulting from focused stimulation of otolith irregular fibers (using a rapid head jerk) remains unknown. Understanding central activity patterns after irregular fiber stimulation lays the groundwork for future studies to investigate central neuronal changes underlying vestibular disorders. Previous studies have used galvanic stimulation (GVS) to activate the vestibular nerve and assess central activity [20]. This low frequency stimulus is purported to preferentially activate irregular fibers throughout the entire vestibular nerve [21]. Our lab has used manganese-enhanced magnetic resonance imaging (MEMRI), a method which takes advantage of the paramagnetic and activity dependent properties of manganese [22] to demonstrate changes in neuronal activity following jerk stimulation [23]. While MEMRI allows for *in vivo* longitudinal comparisons across activated brain regions, individual activated neurons cannot be discerned. To evaluate the distribution and localization of activated neurons, we measured the production of c-Fos in VNC and VeCb subdivisions following jerk stimulation to activate the otolith organs in rats. Production of the immediate early gene, *c-Fos*, has long been used as the gold standard for identifying activated neurons in the brain [24]. Our findings lay the groundwork for future studies to investigate changes in the neuronal function (e.g., activity level and synchrony) following vestibular trauma.

## Materials and Methods

All procedures were approved by the Institutional Animal Care and Use Committee (IACUC) at the University of Michigan (UofM). All rats were individually housed and maintained at 77°F with a 12 – hour light/dark cycle (lights on at 7 AM). Standard housing conditions with free access to normal rat chow and tap water were provided at the UofM AAALAC-accredited animal facility under the supervision of the Unit for Laboratory Animal Medicine (ULAM). After arrival at the UofM ULAM facility, animals were allowed to acclimate for at least 24 hours before undergoing surgical procedures and the eight rats were divided into two groups: a sham group (n=4) and a stimulation group (n=4). All rats underwent surgery (described below), to place a head bolt on the skull, which permitted attachment of the head to a shaker. Ten days after surgery, rats were perfused transcardially and tissue was collected to perform c-Fos immunohistochemistry in the VNC and VeCb of the brain.

### Surgical Placement of Head Bolt

Adult male Sprague-Dawley rats (n=8) were positioned on a stereotaxic frame under ketamine and xylazine anesthesia (ketamine/xylazine 75/8 mg/kg i.p.), and a dorso-cranial midline incision was made to expose bregma and lambda. Once the skull was leveled, two jeweler screws were placed into the skull and a custom head bolt was bonded to the skull with C&B Metabond (Parkell, Edgewood, NY) cement. Dental acrylic was used to secure the head bolt and jeweler screws to the skull (Lang Dental Manufacturing Co., Inc. Wheeling, IL). Following the surgery, animals were allowed to recover for at least 10 days.

### Jerk Stimulation

Ten days post-surgery, Sprague-Dawley rats (n=8) were anesthetized (ketamine/xylazine cocktail – 75/8 mg/kg i.m.) and attached to a mechanical shaker arm via the head bolt (Figure 1A) to transmit the motion of the shaker to the rat’s head. Each animal was positioned to deliver a linear acceleration stimulus in the naso-occipital plane._An accelerometer mounted on the shaker arm confirmed the delivery of a 3,900 g/s jerk stimulus (Figure 1A). Stimulated animals were exposed to 15 trials divided into three blocks (Figure 1B). Each block consisted of 2,000 jerks (6,000 jerks total). One trial equaled 400 jerks (200 jerks up and 200 jerks down). Each block was followed by a 10-minute quiescent interval. Sham animals were also placed in the supine position on the shaker platform, but no jerk stimulus was delivered.

**Figure 1:**
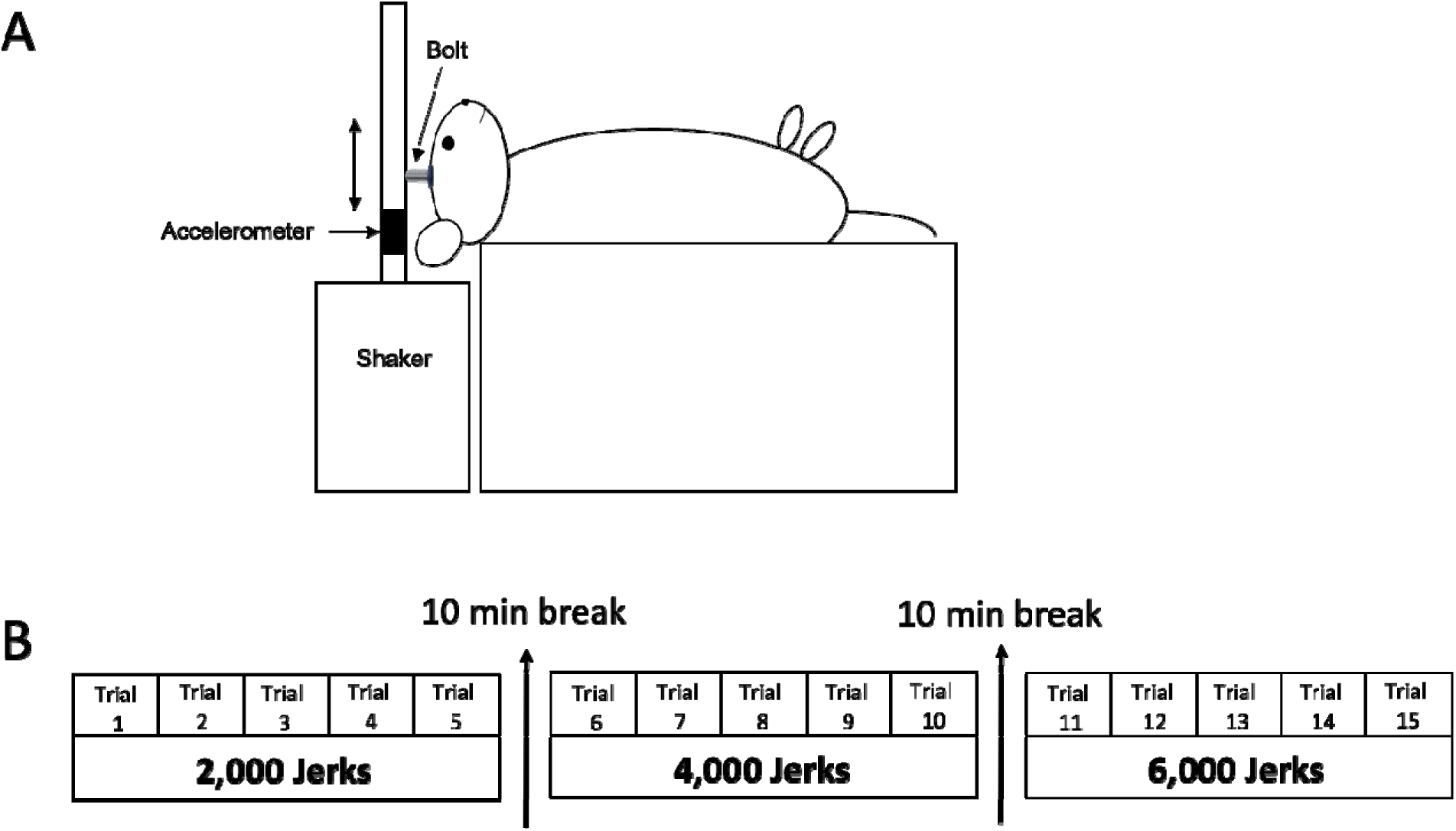
Jerk stimulation paradigm. Rats were placed in the supine position on the shaker platform (A). A head bolt affixed to the skull was attached to the shaker arm in the orientation as shown. Jerk stimuli were applied in the naso-occipital plane (double headed arrows). The jerk stimulation paradigm (B) included a 10-min quiescent interval between each block. Each block consisted of five trials with 400 bidirectional jerks delivered in each trial (i.e., 2,000 jerks per block). The bolded number of jerks at the bottom represents the cumulative number of bidirectional jerks delivered by the end of each block.

### Vestibular Short-Latency Evoked Potential (VsEP)

To record VsEPs, needle electrodes were placed at the vertex (recording), mandible (reference), and hip (ground) of each rat (n=4). Signals from the electrodes were amplified, filtered and digitized using a CED Power 1401 data acquisition system (Cambridge Electron Design LtD Cambridge, MA) at 20 kHz using Spike2 software (Cambridge Electron Design LtD, Cambridge, MA, v7.12). Signals were analyzed using custom MATLAB (MatWorks, Natick, R2019a) scripts, which filtered and averaged all 400 voltage responses in each trial to generate an average VsEP. The signals were synchronized using the stimulus onset as a trigger.

### Transcardial Perfusion and Tissue Collection

Ninety minutes following the onset of jerk or sham stimulation, the anesthetized animals were transcardially perfused with a saline rinse followed by freshly depolymerized 4% paraformaldehyde in PBS. Brains were collected and post-fixed in paraformaldehyde for one hour. Following fixation, brains were placed in 30% sucrose solution overnight at 4°C. Coronal sections (40 µm) through the VNC and cerebellum were serially cut on a freezing sliding microtome (American Optical, Buffalo NY). Sections containing subdivisions of the VNC (Figure 2) – superior vestibular nucleus (SuVe), lateral vestibular nucleus (LVe), medial vestibular nucleus, magnocellular (MVeMC), medial vestibular nucleus, parvocellular (MVePC), and spinal vestibular nucleus (SpVe) - and regions of the vestibulocerebellum (Figure 3) – nodulus (Nod), uvula (Uv), paraflocculus (PFl), and flocculus (Fl) were collected and used for c-Fos immunohistochemistry. Rostral, mid rostro-caudal, and caudal regions of each nucleus were defined and assessed.

**Figure 2.**
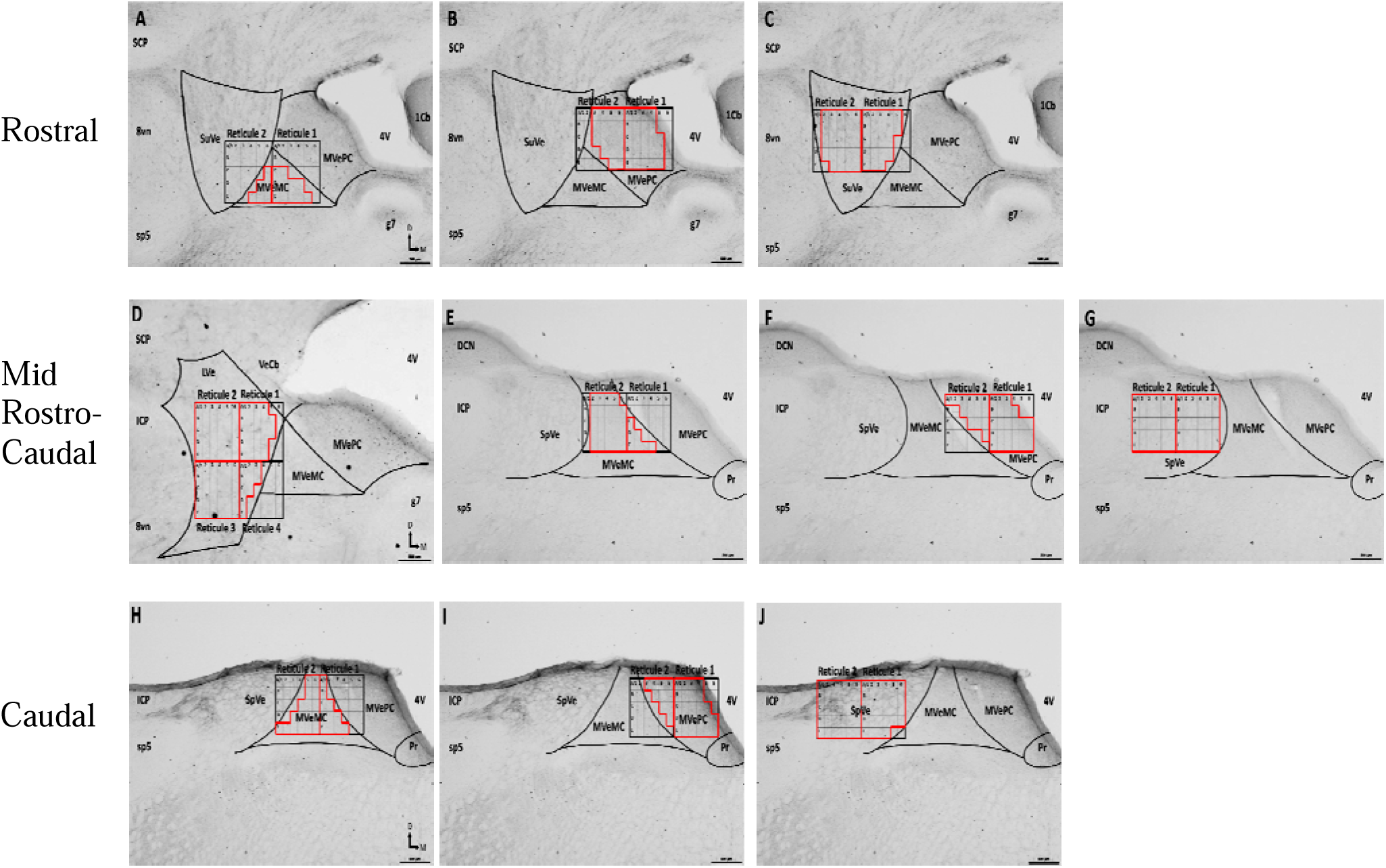
The specific neuroanatomic areas from which c-Fos immunolabeled nuclei counts were performed include rostral (A-C), mid rostro-caudal (D-G), and caudal (H-J) subnuclei of the vestibular nuclear complex (VNC). Black square represents superimposed reticule on photomicrographs; red lines represent specific grid squares (100 um) unique to particular Rols; letters A-E specify rows and 1-6 define the columns of the reticule with unique combinations used to identify ROIs for each subdivision. **Rostral** VNC: MVeMC (A); MVePC (B); SuVe (C); **Mid Rostro-Caudal** VNC: LVe (D); MVeMO (E); MVePC (F); Sp Ve (G); **Caudal** VNC: MVeMC (H); MVePC (I); SpVe (J). **Abbreviations**: 1Cb: first cerebellar lobule; 4V, fourth ventricle; 8vn: vestibulocochlear nerve; DCN: dorsal cochlear nucleus; ICP: inferior cerebellar peduncle; LVe: lateral vestibular nucleus; MVeMC: medial vestibular nucleus - magnocellular portion; MVePC: medial vestibular nucleus - parvocellular portion; Pr: prepositus nucleus; SCP: superior cerebellar peduncle; sp: spinal tract of the trigeminal nerve, Sp Ve: spinal vestibular nucleus, SuVe: superior vestibular nucleus; VeCb: vestibulocerebellum. Reticule - the measurement grid overlaying the tissue (600 µm x 500 µm) used to count Fos labeled nuclei in specific brain regions. One grid box within the reticule =10,000 µm^2^.

**Figure 3.**
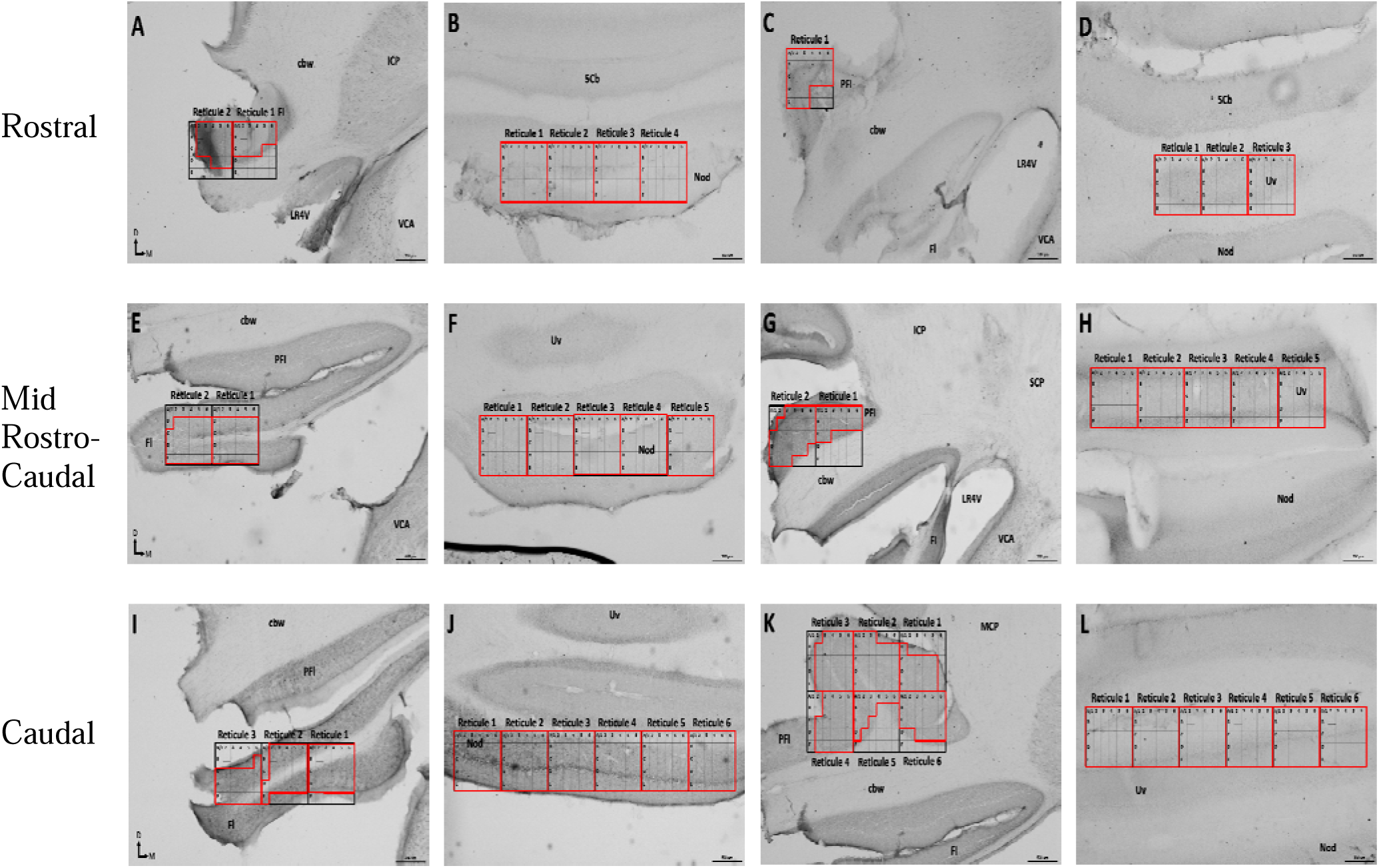
The specific neuroanatomic areas from which c-Fos immunolabeled nuclei counts were performed from r stral (A-D), mid rostro-caudal (E-H), and caudal (I-L) subnuclei of the vestibulocerebellum (VeCB). Black square represents superimposed reticule on photomicrographs, red lines represent specific grid squares unique to particular ROls, letters A-E specify rows and 1-6 define the columns of the reticule with unique combinations used to identify ROIs. **Rostral** VeCb: Fl (A); Nod (B); PFI (C); Uv (D); **Mid Rostro-Caudal** VeCb: Fl (E); Nod (F); PFl (G); Uv (H); **Caudal** VeCb: Fl (I); Nod (J); PEI (K); Uv (L). **Abbreviations**: 5Cb: fifth cerebellar lobule, cbw: cerebellar white matter, FL: flocculus, ICP: inferior cerebellar peduncle, LR4V: left recess of the 4th ventricle, MCP: middle cerebellar peduncle, Nod: Nodulus, PFl: paraflocculus, SCP: superior cerebellar peduncle, Uv: Uvula, VCA: anterior ventral cochlear nucleus. Reticule - the measurement grid overlaying the tissue (600 µm x 500 µm) used to count Fos labeled nuclei in specific brain regions. One grid box within the reticule =10,000 µm^2^.

### c-Fos Immunohistochemistry

Sections were rinsed three times for 15 minutes each in phosphate buffered saline (PBS) and then incubated in a blocking solution consisting of 3% normal goat serum (NGS), 0.5% Triton X-100 (Sigma Aldrich, St. Louis, MO), and PBS for one hour at room temperature on a rotator. Sections were then incubated for 24 hours with a mouse monoclonal c-Fos antibody (1:300; sc-166940, Santa Cruz Biotechnology, Santa Cruz, CA) in PBS and 1% NGS at 4°C on a rotator. After rinsing three times in PBS (15 minutes/rinse), the sections were incubated with a biotinylated secondary antibody (1:200; BA-2000-1.5; Vector Laboratories, Burlingame, CA) in PBS for 1 hour at room temperature on a rotator. Three 15-minute rinses in PBS were followed by a one hour room temperature incubation in an avidin-biotin peroxidase PBS solution (Vector Laboratories, Burlingame, CA). Finally, c-Fos was visualized using diaminobenzidine as the chromogen.

## Analyses

### VsEPs

In all stimulated animals, VsEP responses to jerks were recorded. The amplitude, latency, and duration of three VsEP peaks (P1, P2, and P3) were analyzed to confirm the effectiveness of the jerk stimuli to activate vestibular afferents.

### c-Fos Quantification

Immunolabeled sections from the rostral, mid rostro-caudal, and caudal regions of the VNC and VeCb were examined. Neuronal activity across brain regions was determined by counting the number of c-Fos-immunoreactive nuclei within specific VNC and VeCb subdivisions. With the aid of the Paxinos and Watson Rat Atlas [25], neuroanatomical landmarks were used to define the subdivisions, and images containing each subdivision were acquired (Leica DM4500 microscope; Figures 2-3). A 600 x 500 µm reticule was placed in each subdivision and specific rules applied to track the counts (NIS Elements software, Nikon). The reticule (6 columns (1-6) and 5 rows (A-E) with each square measuring 100 µm x 100 µm) was superimposed on each subdivision. The number of c-Fos immunolabeled nuclei were counted from images obtained at a magnification of 200x (Figures 2-3). To avoid double counting, c-Fos immunolabeled nuclei that touched a gridline were not counted. Counts from each subdivision were normalized to 1000 µm^2^ and reported per mm^2^. To determine the effect of stimulation and rostro-caudal level for each subdivision, a repeated measures ANOVA (p ≤ 0.05) was used followed by a Scheffe test for *post hoc* comparisons (Statview 5.0).

### Vestibular Nuclear Complex Boundaries and Rules for c-Fos Counting

The most *medial* portion of the VNC is bordered by the lateral wall of the 4^th^ ventricle, rostro-caudally, while the *lateral* boundaries are defined by the medial inferior cerebellar peduncle, rostrally. Caudally, the lateral boundaries are defined by the spinal tract of the 5^th^ cranial nerve and nucleus X. The *dorsal* and *ventral* boundaries of the VNC are defined by the floor of the 4^th^ ventricle and either the genu of the 7^th^ nerve rostrally or, for more caudal subdivisions, the floor of the 4^th^ ventricle and the prepositus nucleus. For the current study, the most caudal VNC subdivisions begin in the 40 µm coronal section immediately caudal to the final section containing any of the dorsal cochlear nucleus. The regions of interest were defined in each section and specific rules for identifying and counting the number of c-Fos labeled nuclei in each subdivision of the VNC were specified (Figure 2 and Table 1).

**Table 1.**
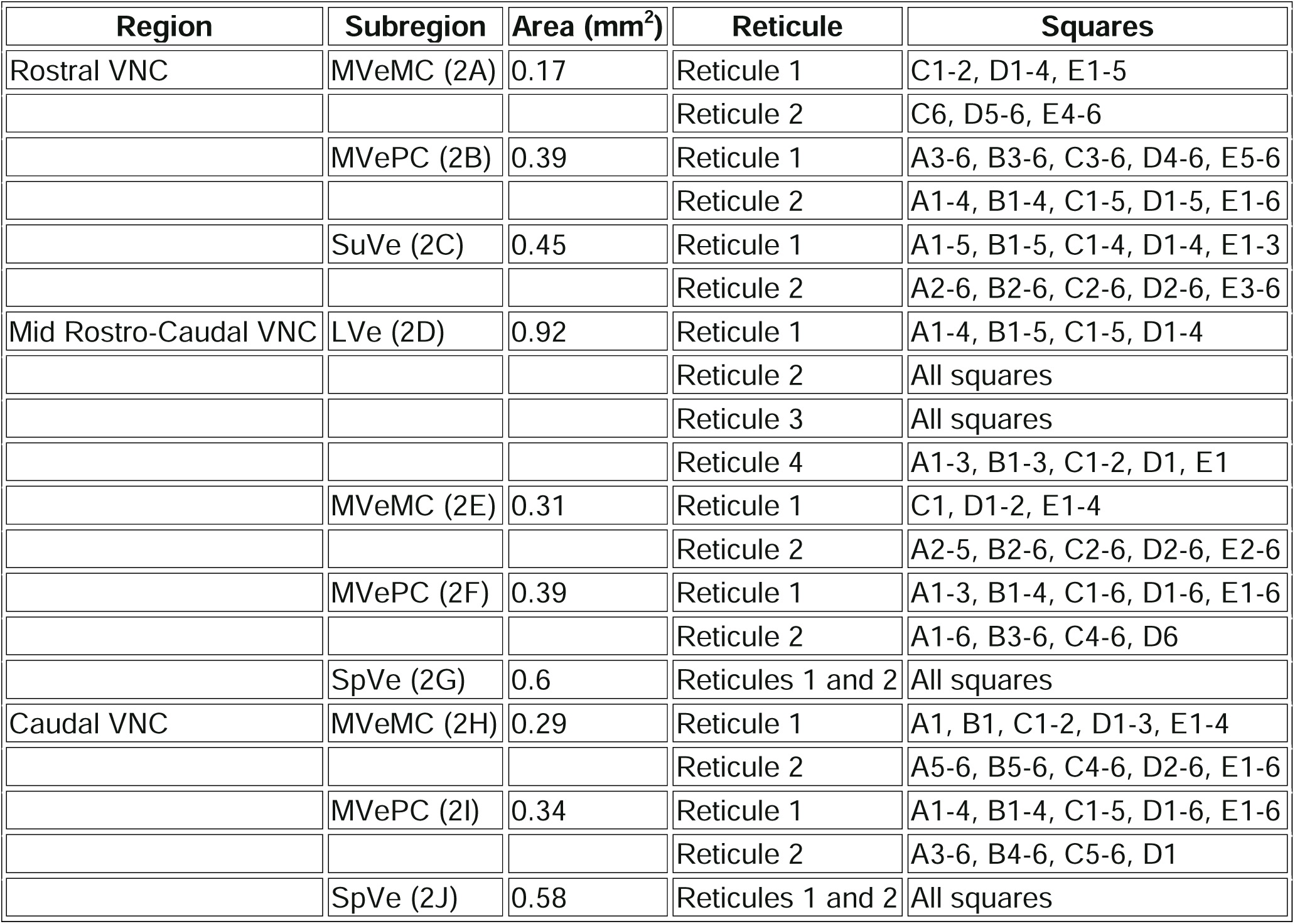
The table provides an overview of the rostro-caudal areas assessed across the VNC and organized by subregion, reticule area, reticule number, and the grid squares included for each region (depicted in Figure 2). Reticule – the measurement grid overlaying the tissue (600 µm x 500 µm) used to count the Fos nuclei in specific brain regions.

<u>The **rostral VNC**</u> contained the SuVe, MVePC, and MVeMC. The number of grid squares used for c-Fos counts was determined by the size and shape of each subdivision (Figure 2 A-C). The grid squares counted for the SuVe were: reticule 1: A1-5, B1-5, C1-4, D1-4, E1-3; reticule 2: A2-6, B2-6, C2-6, D2-6, and E3-6. The MVeMC is overlayed by columns 1-3 in row C, columns 1-4 in row D, and columns 1-5 in row E. For the MVePC, columns 1-4 were counted for rows A and B, columns 1-3 in rows C-E. The **<u>mid rostro-caudal VNC</u>** contained the LVe, both subdivisions of the MVe, and SpVe (Figure 2 D-G). The columns counted for the LVe are 1-5 in rows A-E. Columns 2-6 were counted for the MVeMC. In the MVePC, the columns counted were 1-3 in rows A and B, columns 1-4 for row C, and columns 1-5 rows D and E. All columns in the reticule were counted for the SpVe. The regions classified as part of the **<u>caudal VNC</u>** are both subdivisions of the MVe, and the SpVe (Figure 2 H-J). The MVeMC consisted of grid columns 1-2 in rows A-E. The columns counted for the MVePC are columns 3-5 in rows A-C, columns 3-6 in rows D-E. All columns in the reticule were counted for the SpVe (Figure 2 and Table 1).

### Vestibulocerebellum Boundaries and Rules for c-Fos Counting

The caudal vermis (nodulus, uvula) and the tonsil of the cerebellum (paraflocculus and flocculus) constitute the VeCb. Each of these nuclei extend the entire rostro-caudal length of the VeCb. The regions of interest were defined and specific rules for identifying and counting the number of cFos labeled nuclei in each subdivision of the vestibulocerebellum were specified (Figure 3 and Table 2).

**Table 2.** The table provides an overview of the rostro-caudal areas assessed across the VeCb and organized by subregion, reticule area, reticule number, and the grid squares included for each region (depicted in Figure 3). Reticule – the measurement grid overlaying the tissue (600 µm x 500 µm) used to count Fos labeled nuclei in specific brain regions.

| Region | Subregion | Area (mm<sup>2</sup>) | Reticule | Squares |
| --- | --- | --- | --- | --- |
| Rostral VeCb | FI (3A) | 0.34 | Reticule 1 | A1-6, B1-6, C1-4 |
|  |  |  | Reticule 2 | A2-6, B2-6, C2-6, D4-6 |
|  | Nod (3B) | 1.2 | Reticule 1-4 | All squares |
|  | PFI (3C) | 0.24 | Reticule 1 | A1-6, B1-6, C1-6, D1-3, E1-3 |
|  | Uv (3D) | 0.9 | Reticule 1-3 | All squares |
| Mid Rostro-Caudal VeCb | FI (3E) | 0.47 | Reticule 1 | A1-6, B1-6, C1-6, D1-6, E1-6 |
|  |  |  | Reticule 2 | B2-6, C1-6, D1-6, E1-6 |
|  | Nod (3F) | 1.5 | Reticule 1-5 | All squares |
|  | PFI (3G) | 0.37 | Reticule 1 | A1-6, B1-6, C1-2 |
|  |  |  | Reticule 2 | A3-6, B2-6, C1-6, D1-5, E1-3 |
|  | Uv (3H) | 1.5 | Reticule 1-5 | All squares |
| Caudal VeCb | FI (3I) | 0.65 | Reticule 1 | A1-6, B1-6, C1-6, D3-6 |
|  |  |  | Reticule 2 | A2-6, B2-6, C2-6, D1-6, E1 |
|  |  |  | Reticule 3 | B6, C1-6, D1-6, E1-6 |
|  | Nod (3J) | 1.8 | Reticule 1-6 | All squares |
|  | PFI (3K) | 1.24 | Reticule 1 | B1, C1-5, D1-5, E1-5 |
|  |  |  | Reticule 2 | A1-3, B1-6, C1-6, D1-6, E1-6 |
|  |  |  | Reticule 3 | A3-6, B2-6, C2-6, D2-6, E2-6 |
|  |  |  | Reticule 4 | A3-6, B3-6, C2-6, D2-6, E2-6 |
|  |  |  | Reticule 5 | A1-6, B1-3, C1-2, D1 |
|  |  |  | Reticule 6 | A1-6, B1-6, C1-6, D3-6 |
|  | Uv (3L) | 1.8 | Reticule 1-6 | All squares |

In the **<u>rostral VeCb</u>** (Figure 3 A-D), the grid squares defined for the flocculus were columns 2-6 in rows A and B, columns 3-6 in rows C and D, and columns 4-6 in row E. The grid columns counted for the paraflocculus are 1-4 in row A, 1-5 in rows B and C, and 1-6 in row D. In the **<u>mid rostro-caudal</u>** (Figure 3 E-H) **<u>and caudal</u>** (Figure 3 I-L) VeCb, the grid columns defined for the flocculus are 1-6 in rows A and B, columns 2-6 in rows C and D, and columns 3-6 in row E. The grid squares counted for the paraflocculus are columns 1-4 in row A, 1-5 in rows B and C, and 1-6 in row D. The nodulus and uvula have all grid squares counted throughout the rostro-caudal extent of the VeCb (Figure 3 and Table 2).

## Results

### Jerk stimulation produced comparable VsEPs across animals

For each stimulated rat, VsEP P1, P2 and P3 responses, amplitudes and latencies, were measured to confirm that stimuli were comparable across animals (Figure 4). Responses were consistent across animals with an average P1 amplitude of 0.053 µv ± 0.017 µv and average latency of 0.521 msec ± 0.017 msec (Figure. 4) and an average P2 amplitude of 0.159 µv ± 0.037 µv with a latency of 1.150 msec ± 0.037 msec. These VsEP results show that the jerk stimuli were capable of producing synchronous neuronal activity in the vestibular nerve and brain. Immunolabeling for c-Fos was used to evaluate the number and distribution of active neurons within the VNC and VeCb.

**Figure 4.**
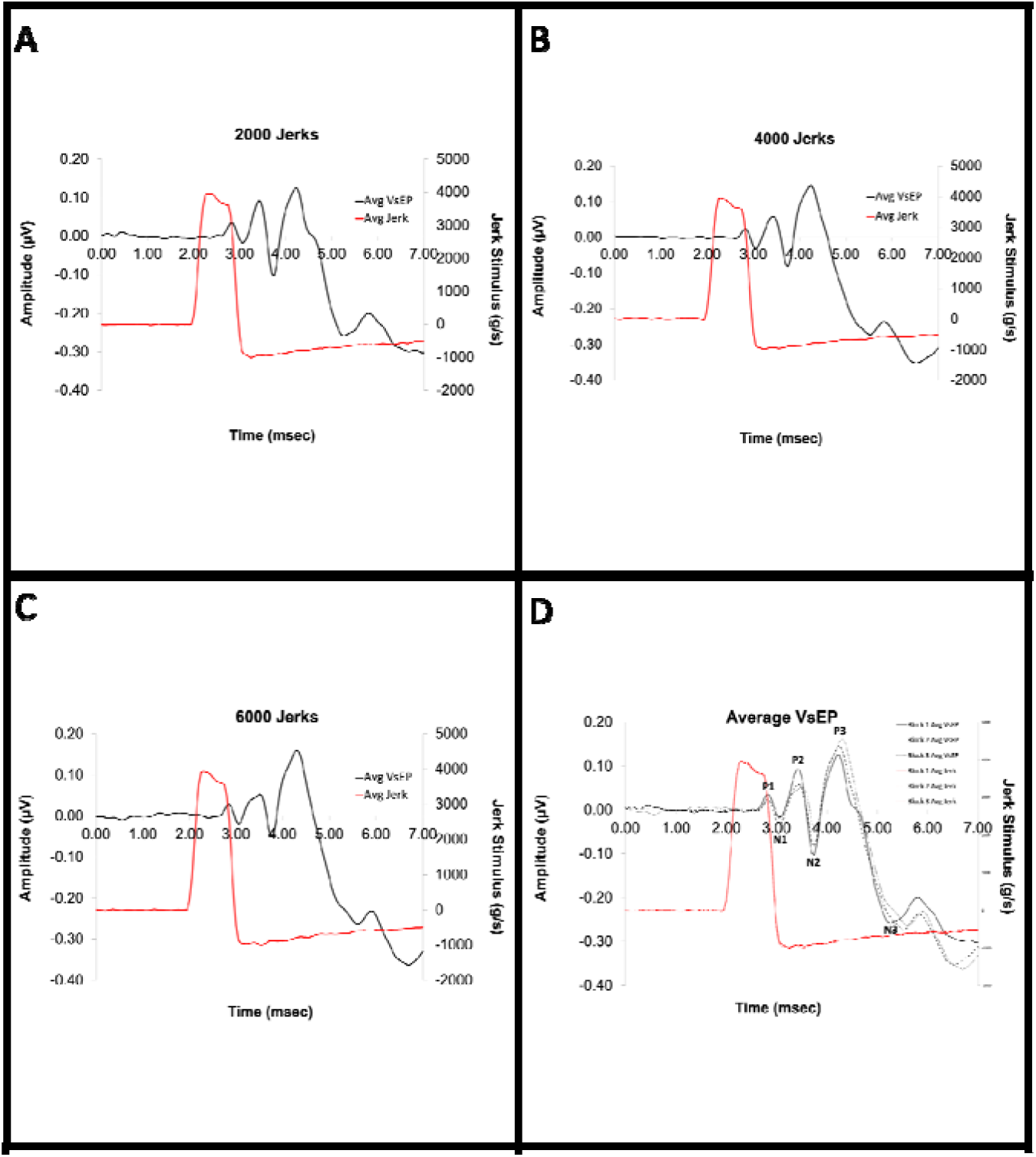
Individual VsEP waveforms represent averaged responses after 2,000 (A), 4,000 (B), and 6,000 (C) jerks. Waveforms for 2,000, 4,000, and 6,000 jerks are superimposed (D). Jerk stimulus is 3,900 g/s (red trace A-D). Y axis left (A-D) - VsEP amplitude; Y axis right (A-D) - jerk stimulus amplitude. P is the positive peak of the wave and N is the negative peak of the wave. (n = 4 rats)

### c-Fos is produced throughout the rostro-caudal extent of the VNC and VeCb in sham rats

In naïve rats not subjected to jerk stimulation (control rats), labeling of c-Fos positive nuclei was identified in both the VNC and VeCb of control rats (Figure 5). The average number of c-Fos positive nuclei was counted within each subdivision of the VNC and VeCb (Table 3, Figure 6 - 7). The results show that central VNC neurons were active even without jerk stimulation. There were significantly more c-Fos positive nuclei found in mid rostro-caudal (p ≤ 0.003) and caudal VNC and VeCb, when compared to rostral subdivisions within these regions (Table 3, Figure 2, Figure 6 - 7). Surprisingly, in the caudal VNC, the MVePC had the greatest number of c-Fos positive nuclei while the SpVe had the fewest c-Fos positive nuclei (Table 3, Figure 6). For the VeCb, the nodulus consistently had the greatest number of c-Fos positive nuclei throughout the rostro-caudal extent of the nucleus (Table 3; Figure 7).

**Figure 5.**
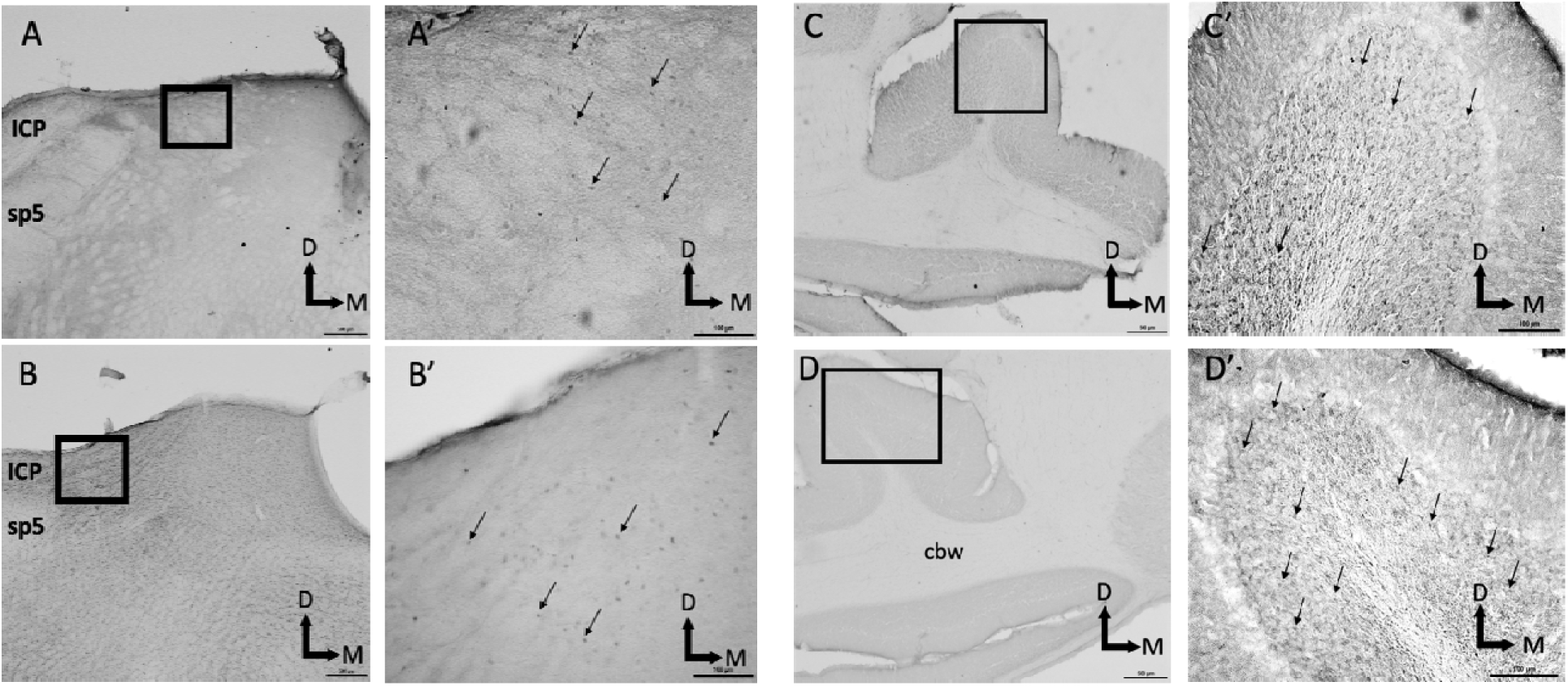
The representation of c-Fos immunolabeled nuclei in the spinal vestibular nucleus in the VNC (A,B) and the paraflocculus of the VeCb (C,D) of control (A and C) and stimulated (B and D) animals. D - dorsal; M – medial; Arrows indicate Fos immuno-positive nuclei. Regions in A’, B’, C’, and D’ represent enlargements of regions in black boxes shown in A, B, C and D.

**Figure 6.**
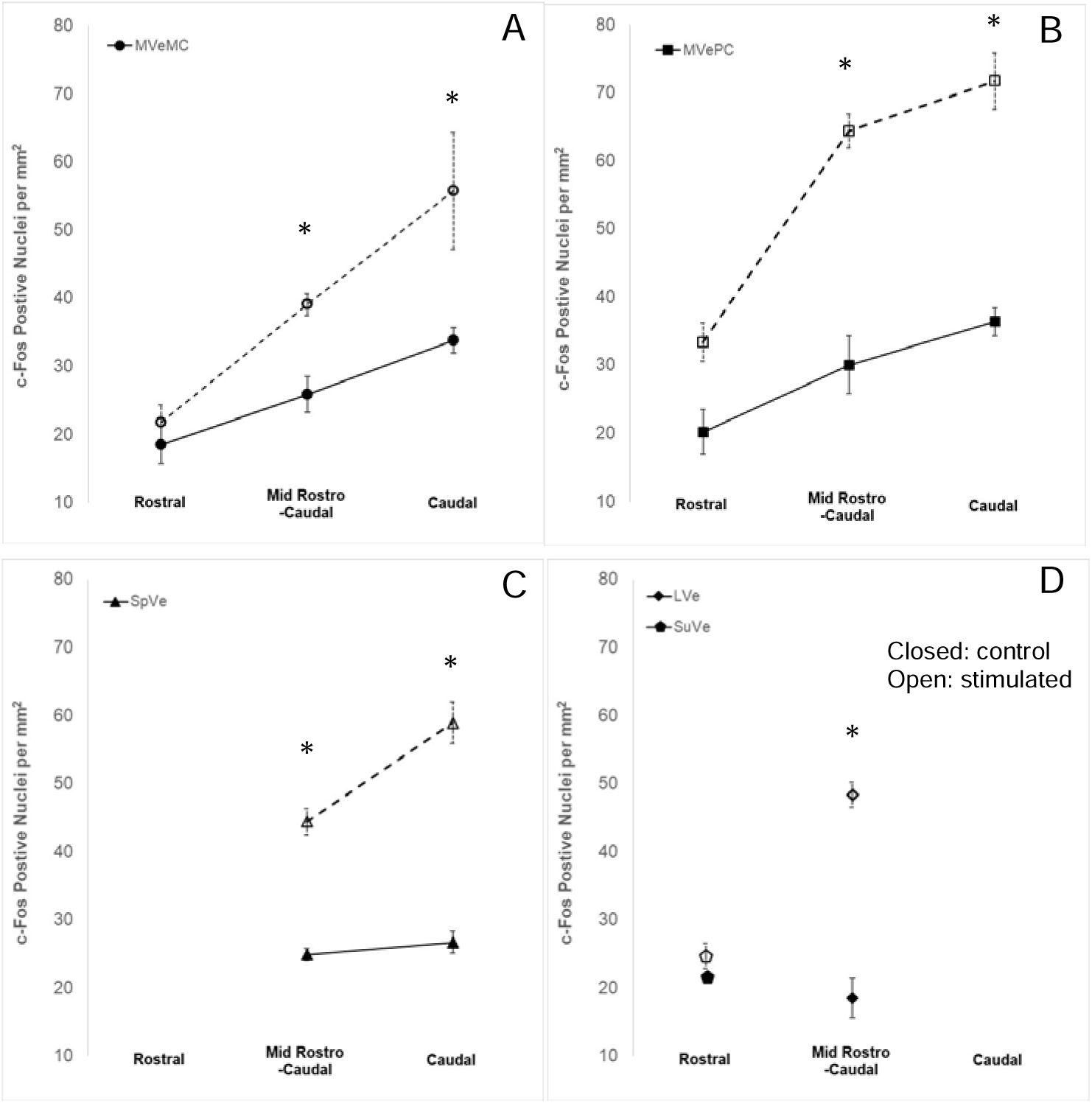
c-Fos production in rostral, mid rostro-caudal, and caudal VNC subdivisions of control (closed symbols) and stimulated (open symbols) animals. Number of normalized c-Fos immunoreactive nuclei counted reported as the mean per mm^2^ ± SEM for the MVeMC (A), MVePC (B), SpVe (C), SuVe, and LVe (D). Asterisks indicate significant difference between control and stimulated animals. Abbreviations: MVeMC - medial vestibular nucleus, magnocellular division; MVePC - medial vestibular nucleus, parvocellular division; SpVe - spinal vestibular nucleus; SuVe - superior vestibular nucleus; error bars equal SEM; (n = 4 rats per group)

**Figure 7.**
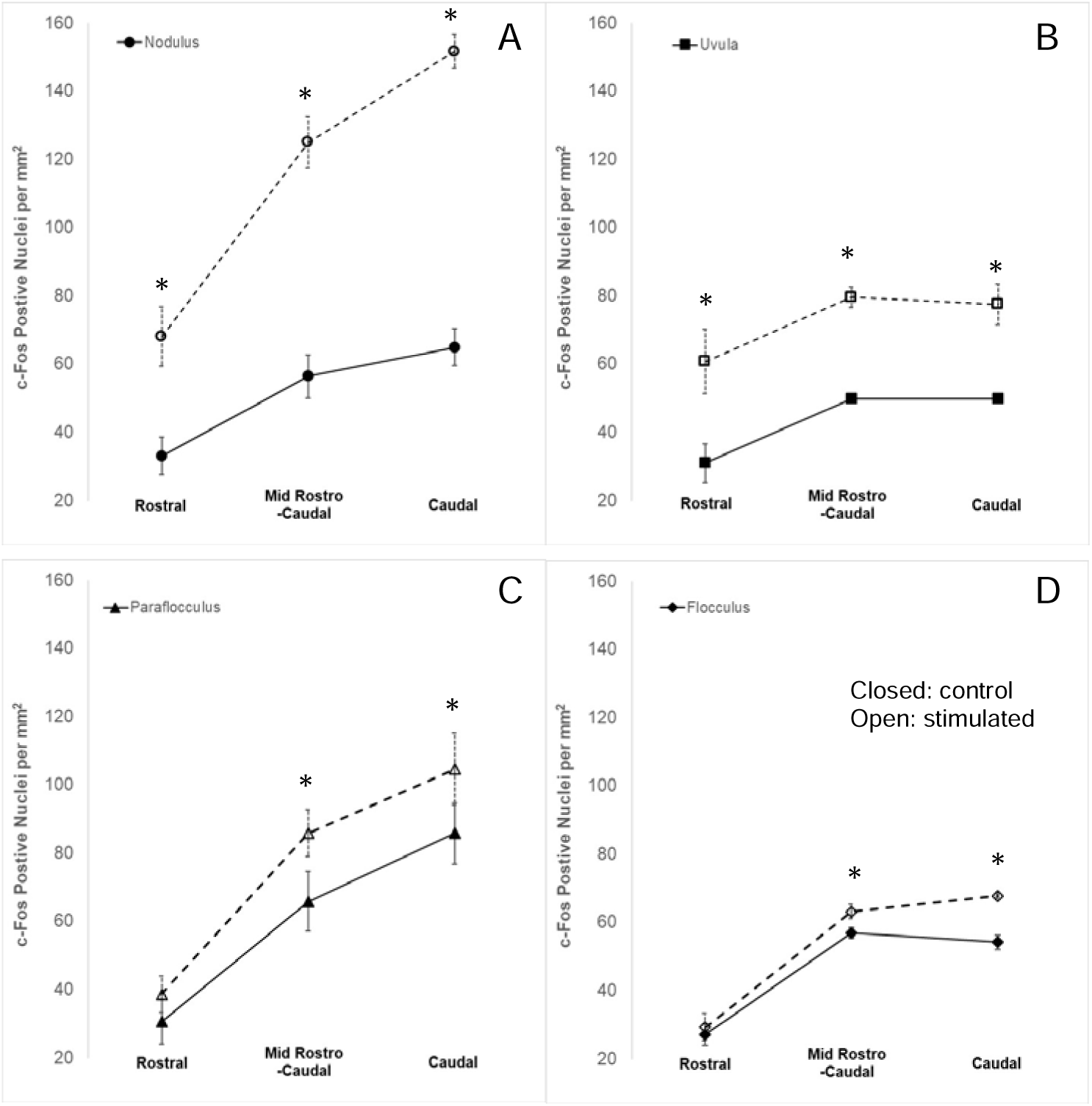
c-Fos production in rostral, mid rostro-caudal, and caudal VeCB subnuclei of control (closed symbols) and stimulated (open symbols) animals. Number of c-Fos immunoreactive nuclei counted reported as the mean per mm^2^ ± SEM for the Nod (A), Uv (B), paraflocculus (C), and flocculus (D). Asterisks indicate significant difference between control and stimulated animals. Abbreviations: Nod - nodulus; Uv - uvula; PFL - paraflocculus; FL – flocculus; error bars equal SEM; (n = 4 rats per group)

**Table 3.** Comparison of the mean number of c-Fos immunolabeled neurons per mm^2^ in VNC and VeCb subdivisions of control and jerk stimulated groups. Values are mean ± SEM (n = 4 rats per group)

| Subnucleus | Control | Stimulated |
| --- | --- | --- |
| **VNC** |  |  |
| **Rostral** |  |  |
| MVeMC | 18.43 ± 2.77 | 21.78 ± 2.57 |
| MVePC | 20.16 ± 3.31 | 33.36 ± 2.83 |
| SuVe | 21.42 ± 4.24 | 24.71 ± 1.86 |
| **Mid Rostro-Caudal** |  |  |
| LVe | 18.54 ± 2.92 | 48.38 ± 1.83 |
| MVeMC | 25.84 ± 2.64 | 39.12 ± 1.60 |
| MVePC | 30.03 ± 4.25 | 64.40 ± 2.48 |
| SpVe | 24.92 ± 0.88 | 44.42 ± 1.90 |
| **Caudal** |  |  |
| MVeMC | 33.85 ± 1.86 | 55.79 ± 8.59 |
| MVePC | 36.40 ± 2.03 | 71.73 ± 4.15 |
| SpVe | 26.71 ± 1.60 | 58.97 ± 3.02 |
| **VeCb** |  |  |
| **Rostral** |  |  |
| Nodulus | 33.19 ± 5.52 | 68.09 ± 8.79 |
| Uvula | 30.99 ± 5.63 | 60.76 ± 9.45 |
| Paraflocculus | 30.64 ± 6.64 | 38.71 ± 5.30 |
| Flocculus | 27.17 ± 3.14 | 29.33 ± 4.02 |
| **Mid Rostro-Caudal** |  |  |
| Nodulus | 56.39 ± 6.30 | 125.01 ± 7.55 |
| Uvula | 49.86 ± 1.40 | 79.66 ± 3.00 |
| Paraflocculus | 65.84 ± 8.77 | 85.84 ± 6.92 |
| Flocculus | 56.94 ± 1.63 | 63.17 ± 2.11 |
| **Caudal** |  |  |
| Nodulus | 64.92 ± 5.42 | 151.61 ± 4.87 |
| Uvula | 49.85 ± 1.40 | 77.48 ± 5.87 |
| Paraflocculus | 85.80 ± 8.89 | 104.65 ± 10.55 |
| Flocculus | 54.26 ± 2.12 | 67.81 ± .75 |

### Jerk stimulation does not change the number of c-Fos immunolabeled nuclei in the rostral VNC

The average number of c-Fos positive nuclei were compared in control and stimulated animals. Three rostral VNC subdivisions (MVeMC, MVePC,and SuVe) were evaluated (Table 3, Figure 6). No significant differences were found in the number of immunolabeled c-Fos positive nuclei after stimulation compared to sham controls within each of these rostral subdivisions (Figure 6).

### Jerk stimulation increases c-Fos production in mid rostro-caudal and caudal VNC subdivisions

Each mid rostro-caudal and caudal subdivision of the VNC (Figure 6) had significantly increased numbers of c-Fos immunolabeled nuclei following jerk stimulation (p ≤ 0.05). Of the mid rostro-caudal VNC subdivisions the MVePC had the largest number of c-Fos immunolabeled nuclei per mm^2^ after jerk stimulation (Figure 6B). However, the LVe had the largest change, with a 2.6 fold increase in c-Fos labeled nuclei after stimulation (Figure 6D), while the MVePC and SpVe subdivisions had 2.0 - and 1.8 fold increases respectively (Figure 6B, 6C). Neurons in the MVeMC had the smallest fold change after jerk stimulation (1.5 fold). These findings in the mid rostro-caudal VNC were similar to those in the caudal VNC (Figure 6). The increases in number of c-Fos immunolabeled nuclei following jerk stimulation were the smallest in the caudal MVeMC (Figure 6A) when compared to the caudal MVePC (Figure 6B). However, it was the caudal SPVe that showed the largest change with a 2.21 fold increase (Figure 6C) following jerk stimulation.

### Jerk stimulation differentially increases c-Fos production in VeCb subnuclei

In rostral regions of the VeCb only the Nod and Uv have significant increases in c-Fos labeled nuclei after jerk stimulation when compared to sham controls (Figure 7A<u>, 7B)</u>. However, in addition to the Nod and Uv, c-Fos immunolabeled nuclei in the Fl was significantly increased in the mid rostro-caudal region of the VeCB (Figure 7C). Significant increases in c-Fos labeled nuclei in response to jerk stimulation were also observed in all four nuclei in the caudal VeCb (Figure 7), suggesting that linear acceleration significantly contributes to the activity of the VeCb, with caudal regions more affected than rostral regions.

## Discussion

### Neuronal activity in the VNC and VeCb can be evaluated by jerk stimulation

This study provides the first investigation of the impact of jerk stimulation on central neuronal activity across vestibular related nuclei in rats. The study demonstrates that jerk stimulation activates the immediate early gene *c-fos*, in the MVePC, LVe, and SpVe of the VNC and in the Nod, Uv, Fl, and PFl of the VeCb. The largest number of activated cells per unit area were in the MVePC and the Nod. For several subdivisions there was a rostro-caudal c-Fos gradient, with more activated neurons in the mid rostro-caudal and caudal regions of the nucleus (MVeMC, MVePC, SpVe, Nod, PFl, and Fl). We delivered a consistent jerk stimulus of 3,900 g/s which generated reproducible VsEP responses (Figure 4). Although the VsEP wave we identify as P2 is referred to as P1 in some other studies, we attribute the function of this wave to activity in central vestibular pathways based on evidence from lesion studies [7].

### Central neuronal activity is observed in the absence of jerk stimulation

In the current study, c-Fos immunolabeled nuclei were found throughout the VNC and VeCb in the sham group. This result is different from other studies reporting little to no c-Fos labeling in the VNC of sham animals prior to bilateral vestibular nerve stimulation [20, 26]. The type and level of anesthesia in these studies was similar to those used in the current study and therefore should not play a role in the differences observed when comparing the current results to studies reporting negligible c-Fos labeling in the VNC of sham animals. An explanation for this discrepancy may be differences in the concentration, source, and sensitivity of the antibodies. Since c-Fos levels often peak within 90 -180 minutes and can remain detectable for up to 24 hrs, another explanation is that the c-Fos labeling in sham animals may reflect the animals’ baseline vestibular activity *prior* to being placed in the supine position, on the shaker platform. Another explanation for c-Fos labeling in the sham group may be associated with the placement of animals in the supine position on the shaker platform (Figure 1A). Feher (2012) notes that the hair cells within the otolith organs are capable of sensing tonic information in relation to gravity [27]. Therefore, even when the head is at rest, otoconia within the maculae exert a force on the mechanoreceptor which is equal to 9.8 m/sec^2^, due to the gravitational pull of the earth. Given the constant presence of gravitational acceleration the positioning and maintenance of the animals in the supine position for ∼30 minutes, producing the experience of gravity in an unusual direction, may provide a stimulus great enough to engage activity in the otolith organs and account for the c-Fos patterns observed in the sham group. Similar to jerk stimulated animals, sham animals had more c-Fos labeled neurons in caudal subdivisions within the VNC when compared to rostral subdivisions (Table 3). Since otolith projections are more enriched in caudal subdivisions of the VNC, the pattern of c-Fos labeling in shams fits well with the idea of otolith activation.

In addition to otolith projections, the caudal VNC has been linked to the vestibulo-sympathetic reflex (VSR). The VSR provides dynamic modulation of blood pressure, heart, and respiratory rate during changes in posture, contributing to maintenance of appropriate brain perfusion [28]. The VSR pathway sends signals from vestibular end organs to caudal vestibular nuclei. In the current study, placing animals in the supine position could result in hypotensive effects that engage the vestibulo-sympathetic reflex and produce c-Fos labeling. Holstein et al., suggest that neurons distributed throughout the caudal VNC play a role in the vestibulo-sympathetic reflex and are activated following sGVS activation of the vestibular nerve. In those studies, animals were not reported to be in the supine position during vestibular nerve stimulation and very few c-Fos labeled neurons were seen in sham animals [20]. In further support of the VSR contribution to c-Fos labeling in the sham animals, one study have examining the effect of head position on mean arterial pressure found that people in the prone position had significant changes when the head was in the up versus the down position [29]. The results were similar to those observed after bilateral vestibular loss [30].

One method to attempt to reduce vestibular activity prior to jerk stimulation can be restraint of the animals for several hours prior to the jerk stimulus. However, we opted for more natural conditions and did not want to add restraint stress as a variable. In the future, the c-Fos levels found in sham animals should be further investigated, comparing several stimulation positions and techniques (e.g., prone versus supine and jerk versus sGVS). Nonetheless, the effects of gravity and position would have been present in all test animals prior to and during jerk stimulation. Therefore, since increases in c-Fos labeling were observed above that in the controls, the impact of jerk stimulation on c-Fos labeling was able to be assessed.

### Jerk stimulation generates differential neuronal activity

In the current study, a type of rapid linear head movement, jerk, was used to stimulate irregular vestibular afferent fibers. The striola of both the saccular and utricular otolith organs are innervated by rapidly adapting irregular vestibular afferent fibers that project to the VNC and VeCb. Previous studies suggest that when the animal is in the supine position and the jerk stimulus is applied in the naso-occipital plane, the saccule is primarily stimulated [6]. However, since utricular and saccular maculae are at right angles to one another, but are not orthogonal to earth’s axis, utricular stimulation cannot be ruled out [31]. It is unlikely the semicircular canals contributed significantly to central activation in this study since rotational head movement (angular acceleration) was negligible.

### Jerk stimulation increases neuronal activity in the VNC

The labeling we observed was bilateral and symmetrical, similar to the bilateral symmetry reported in the numbers of Fos immunoreactive in intact non-deafferented animals [20, 32–34]. Previous studies have reported the production of c-Fos in the MVe, SpVe, and the SuVe using several paradigms to stimulate otolith organs in rodents [32, 35, 36]. Although GVS has been used to assess effects of stimulation of vestibular end organs on neuronal activity in the VNC [20], activity patterns across the VeCb have not been assessed. However, it is unclear which afferent fiber populations were activated by the sGVS stimuli employed in those studies [21]. The jerk stimulation in the current study specifically targeted irregular fibers of the otolith organs [37] sensitive to abrupt linear motion; some dimorphic and likely all afferents with regular resting discharges are not activated by jerk stimuli. Thus, our assessment of activated neurons across subdivisions of both the VNC and the VeCb is limited to those neurons directly or indirectly activated by jerk stimuli. Our analysis of c-Fos immunolabeling reveal that stimulation of irregular fibers leads to the activation of nuclei in the VNC and VeCb. For the VNC significant increases in the number activated neurons were found in three of the four subdivisions assessed, with the SuVe being the exception. Additionally, substantial neuronal activation was noted in the nodulus and uvula throughout all levels of the VeCb. The pattern of c-Fos production observed under our experimental conditions suggests that jerk stimulation activates VNC and VeCb subdivisions involved in spinal modulation of motor responses with less involvement of those related to oculomotor responses. Our findings support previous literature that suggest that in response to jerk stimulation delivered in the naso-occipital plane, the otolith organs are the primary source of afferent stimulation to the VNC and VeCb.

### VNC regions enriched with otolith projections are most activated

The number of jerk-activated neurons was compared across the rostro-caudal extent of the VNC. The most robust immunolabeling for c-Fos was observed in the mid rostro-caudal and caudal VNC subdivisions. In comparison, fewer jerk-activated neurons were present in rostral VNC subdivisions (Figure 6 – 8). This regional specificity potentially reflects differential processing of vestibular information related to linear acceleration. Neurons in the caudal VNC are reported to receive significant projections from afferents originating in otolith organs, including the LVe, MVe, and SpVe [38]. Results from the current study are similar to earlier studies that reported more activated nuclei in the caudal VNC than in the rostral VNC after sinusoidal galvanic stimulation – sGVS. The results are consistent with the fact that the rostral VNC is more concerned with eye movement and orientation in space (ascending projections). The earlier studies attributed the differences in rostral versus caudal neuronal activation to enriched otolith afferent projections to caudal VNC subdivisions [18]. However, unlike these other studies we have found significant labeling in the LVe. Perhaps the stimulation used in previous studies, which is not selective for particular vestibular afferents, inhibits LVe neuronal activity. Our results suggest that this inhibition of the LVe may not be present when a population of irregular fibers are preferentially stimulated and implicate regular vestibular afferents in LVe inhibition (descending projections).

The MVe is divided into parvocellular (MVePC) and magnocelluar (MVeMC) regions. The MVePC serves as a nexus for a wide array of projections to the VNC [39]. Previous studies suggest that the MVePC, particularly the caudal MVePC, receives a predominance of afferent input from the otolith organs, while the MVeMC receives a preponderance of input from the semicircular canals [18, 40]. Our findings are in line with these observations, as the greatest number of jerk-activated neurons were observed in the MVePC. Following GVS, the highest density of c-Fos positive neurons was concentrated in the MVePC with notably sparse immunolabeling the MVeMC. While jerk-activated neurons in the MVePC may be attributed to direct stimulation of otolith irregular afferents, the MVePC also receive ipsilateral projections from regular afferents as well as bilateral commissural fiber projections. This commissural activation consists of a subpopulation of MVe GABAergic neurons activated by excitatory neurons in the contralateral VNC. In the current study, this secondary activation of these inhibitory GABAergic neurons would be indistinguishable from the direct afferent activation of neurons in the MVe.

As one of the subdivisions forming the medial vestibulospinal tract (MVST), the MVe affects motor neurons controlling neck musculature [41]. Our results suggest that jerk stimulation of the otoliths is a potent, activator of neurons in the MVePC and supports a role for these neurons as significant contributors to the MVST, facilitating rapid compensatory adjustments by motor neurons and interneurons to maintain neck stabilization during head movements.

The results suggest that the jerk stimulation pardaigm used in the current study produces more labeled c-Fos nuclei in VNC subdivisions, which may be specifically related to otolith stimulation. Previous rotation studies have compared the effects of otolith stimulation to semicircular canal stimulation and found differences in the pattern of c-Fos labeling. Additional studies are needed to determine whether linear jerk stimulation produces activation patterns that are different from those observed after angular jerk stimulation.

### VeCb subdivisions receiving direct peripheral projections produce the most c-Fos

The number of jerk-activated neurons was compared across the rostro-caudal extent of the VeCb. Immunolabeling for c-Fos was most robust in the mid rostro-caudal and caudal VeCb subdivisions. Fewer jerk-activated neurons were present in rostral VeCb subdivisions (Figure 7). Activity in the nodulus and uvula was significantly increased throughout the VeCb in response to jerk stimulation. The largest number of jerk-activated VeCb neurons was found in the nodulus. Compared to the paraflocculus and flocculus, the nodulus and uvula receive robust afferent projections originating from the saccule [39]. Our results suggest that these projections are distributed rostrocaudally throughout the VeCb (Figure 7) and may explain the robust jerk-stimulated c-Fos labeling observed throughout the nodulus and uvula. Jerk stimulation produced activated neurons in the mid rostrocaudal and caudal flocculus. However, only the most caudal paraflocculus had significant increases in jerk-activated neurons (Figure 7; Table 1). The VeCb sends partially overlapping projections to nuclei of the VNC [18]. These VeCb to VNC projections are primarily inhibitory [42]. The paraflocculus and flocculus contribute to the coordination of eye movements. Primate studies demonstrate that the ventral paraflocculus has effects on both visuomotor integration and motor control while floccular projections play critical roles in ocular motor control, including the modulation of the VOR and smooth-pursuit eye movements [43]. These results align with previous studies demonstrating interactions between the flocculus and the SuVe [44]. In the current study the fewest jerk-activated VNC neurons were observed in the SuVe and MVeMC (Figure 6; Table 3) and may reflect VeCb mediated limited activation of visuomotor and ocular pathways by naso-occipital jerk stimulation.

## Conclusions

Bidirectional jerk stimulation, which selectively activates irregular fibers sensitive to abrupt linear acceleration, provides a reliable method for mapping the connectivity of these fibers with secondary vestibular neurons in the VNC and VeCb. In both sham and jerk-stimulated rats, the otolith organs are presumed to be the primary source influencing the activity patterns of the vestibular nuclei complex (VNC) and vestibulocerebellum (VeCb).

The jerk stimulation paradigm in the current study was limited to one stimulation intensity of 3,900 g/s. To address the sensitivity of VNC and VeCb activation patterns after irregular fiber stimulation, the effect of different simulation intensities on the number of c-Fos labeled neurons should be assessed. While the degree of activity for a single neuron cannot be determined, the extent to which a given subdivision or rostrocaudal region is activated can be examined. Correlations between the jerk intensity, the number of c-Fos labeled nuclei, and the amplitude of VsEP P1-P3, can be analyzed. This would aid in our understanding of the limitations of VNC and VeCb neurons to be recruited across a broad range of stimuli in response to irregular fiber stimulation. As mentioned above, these results should be further explored in future studies with different paradigms that include comparing position (e.g., supine versus prone), acceleration direction (linear versus angular), etc… The results would allow us to learn more about the relationship between gravity and central pathway activation as well as whether any segregation of activation of VNC and VeCb neurons can be discerned anatomically or morphologically between irregular fiber stimulation originating from the otoliths versus semicircular canals.

Jerk stimulation significantly affects specific regions in central vestibular-related nuclei. Specifically, regions in the VNC and VeCb that have enriched projections from otolith organs demonstrate the greatest change in neuronal activation. Ultimately, increased activation of specific central vestibular-related nuclei impacts both motor and sensory pathways allowing adjustments to neck, trunk, and eye movement. Similar to studies using GVS, future studies should address the localization and distribution of neurochemically identified jerk-activated VNC and VeCb neurons and their contributions to motor and sensory pathways. Understanding the identity and origin of first order neurons driving the jerk-activation of VNC and VeCb neurons will further define the role of irregular afferents in normal and pathological vestibular function.

The production of c-Fos is often used as a marker for activity [45–48]. Other studies, with a focus on vestibular compensation, for example, the production of c-Fos indicates a first step in changes in the cell structure or function that occurs as a consequence of a significant stimulus [34, 49]. Some may refer to this as plasticity. The vestibular paradigms we used provided stimulation of a subset of afferents that was outside of what would be the physiological range of a rat. This may result in generating plastic changes throughout the vestibular pathway. Immunolabeling for c-Fos was observed in both the VNC and VeCb and therefore the potential impact of plastic change would vary broadly and be a function of the neurochemistry of the neuron producing the cFos. In summary, these experiments demonstrate that vestibular stimulation of irregular fibers activates, and potentially drives plastic changes in, the VNC and VeCb as well as to the nuclei to which they project

The study demonstrates that otolith irregular afferent stimulation elicits a widespread, but distinct pattern of neuronal activation within the VNC and VeCb. The most significant changes in neuronal activity were localized to regions with enriched direct otolith inputs. These results contribute to our understanding of the central processing of vestibular information and the neural substrates underlying rapid compensatory responses to head movements.

## Abbreviations

PFl: Paraflocculus
Fl: Flocculus
Nod: Nodulus
Uv: Uvula
4V: Fourth ventricle
SuVe: Superior vestibular nucleus
LVe: Lateral vestibular nucleus
MVeMC: Medial vestibular nucleus magnocellular division
MVePC: Medial vestibular nucleus, parvocellular division
SpVe: Spinal vestibular nucleus
VNC: Vestibular nuclear complex
VeCb: Vestibulocerebellum
VsEP: Vestibular short-latency evoked potential
MEMRI: Manganese-enhanced magnetic resonance imaging
GVS: Galvanic stimulation

## Acknowledgements

We thank Helena Bissinger and Hannah Beck for their technical contributions. We also thank the John D. Dingell VAMC, Wayne State University School of Medicine, VA Ann Arbor Healthcare System, and the Mountain Home VA Healthcare System for their ongoing support of this research. This research was supported by grant R01DC018003-S1 of NIH and NIDCD to MK, grant 1I01RX001986-01 of the US Department of Veteran Affairs to AGH and FA, and grant 1I21RX004111-01A1 of the US Department of Veteran Affairs to AGH. The views expressed do not necessarily reflect the official policies of the Department of Health and Human Services, nor does mention of trade names, commercial practices, or organizations imply endorsement by the U.S. Government.

